# Spatially conserved pathoprotein profiling in the human suprachiasmatic nucleus in progressive Alzheimer disease stages

**DOI:** 10.1101/2024.03.07.584000

**Authors:** Gowoon Son, Mihovil Mladinov, Felipe Luiz Pereira, Song Hua Li, Chia-Ling Tu, Grace Judge, Yumi Yang, Claudia Kimie Suemoto, Renata Elaine Paraízo Leite, Vitor Paes, Carlos A. Pasqualucci, Wilson Jacob-Filho, Salvatore Spina, William W. Seeley, Wenhan Chang, Thomas Neylan, Lea T. Grinberg

## Abstract

Individuals with Alzheimer’s Disease (AD) experience circadian rhythm disorder. The circadian rhythm is synchronized by a master clock, the suprachiasmatic nucleus (SCN), which is a tiny hypothalamic nucleus. Little is known about the molecular and pathological changes that occur in the SCN during AD progression. We examined postmortem brains of 12 controls without AD neuropathological changes (Braak stage 0) and 36 subjects at progressive Braak stages (I, II, and VI). To investigate potential AD-specific changes, we measured the neuronal counts of arginine vasopressin (AVP) and vasoactive intestinal peptide (VIP) positive neurons, along with the Braak stages in the SCN. We investigated in adjacent hypothalamic nuclei which are also composed of AVP+ neurons but show more resilience to AD: paraventricular nucleus (PVN) and supraoptic nucleus (SON). To understand the dysregulated proteins associated to AD progression, we performed in-situ proteomics, investigating 57 proteins, including commonly dysregulated in AD, using GeoMx Digital Spatial Profiling (DSP) in the three nuclei (total of 703 area of interests). Neurofibrillary tangles (NFTs) and tau fibrils were found selectively in SCN. We failed to detect NFTs in SON, only a mild dysregulation of p-tau at Braak VI in PVN and SON. Amyloid plaque was absent in the SCN and SON. Additionally, the SCN showed increased glial proteins already at Braak stage I, whereas the level of these proteins sustained in the other nuclei. The SCN is exclusively vulnerable to AD-tau pathology and show immune dysregulation even at Braak I but is protected against amyloid plaque. This finding revealed selectively in amnestic AD, showing more resilience in AD variant. This tau-related molecular dysregulation in the SCN contributes to circadian rhythm disturbances in AD, a phenomenon observed before the onset of cognitive disorder.

## Introduction

The suprachiasmatic nucleus (SCN) in the hypothalamus is known as the central circadian pacemaker of the human brain. The SCN synchronizes the endogenous oscillations of protein expression from individual neurons and glial cells throughout the entire system (1). It maintains the rhythmic balance of cellular circadian and sleep-wake cycles. The SCN transmits local circadian signals to various subcortical areas, facilitating sleep-wake homeostasis through direct synaptic connections (2,3).

Circadian rhythm disorders are studied as an early symptom in patients with neurodegenerative diseases, characterized by abnormal protein aggregation and fundamental molecular abnormalities. Patients with Alzheimer’s disease (AD) particularly struggle with disrupted daily activity and rest cycles(4). Human studies suggest that the presence of amyloid and tau in patient samples is linked to disruption in physiological rhythmic processes, such as body temperature regulation, hormone balance, and the sleep/wake cycle (5–8). Genetically modified animal models that overexpress human amyloid and tau proteins have suggested a link between the accumulation of these proteins and circadian abnormalities (9). In addition to the amyloid and tau pathology, the dysregulation of inflammatory glial populations and the expression of molecules involved in protein homeostasis have been suggested as contributing factors to amyloid and tau accumulation and circadian dysregulation (10–13). However, previous neuropathological examinations have primarily focused on the hippocampal formation and cortical areas, leaving the impact on the SCN underexplored (14). The neuropathological examination in human SCN is primarily focused on comparing severe AD cases with controls (14). This highlights the need to warrant neuropathological investigations to the early stages of AD. Additionally, none of the previous studies have elucidated the potential mechanisms responsible for the accumulation of pathological proteins in the human SCN.

Therefore, we verified the inclination for AD neuropathological alterations in human SCN and investigated the molecular susceptibility of the SCN during the AD progression. The human SCN, a tiny nucleus of 50,000 neurons (20,000 neurons in mouse) located in one hemisphere of the anterior hypothalamus, consistently shows spatially conserved expression of neuromodulatory cells (15–18). Of the 10 types of neuromodulatory cells present in the SCN, arginine vasopressin (AVP) and vasoactive intestinal peptide (VIP) are acknowledged as the primary regulators of circadian rhythms. AVP-positive (AVP+) and VIP-positive (VIP+) neurons constitute 30% of the SCN neurons and govern cellular circadian rhythms with distinctive spatial distribution; AVP neurons predominantly populate the shell, while VIP neurons are concentrated in the core of the SCN (17). The two primary neuromodulatory neurons transmit cellular information from the SCN to other subcortical nuclei. AVP cells are recognized for transferring the efferent circadian signal from the SCN to the PVN and other hypothalamic nuclei (19,20). Conversely, VIP cells process the afferent signals from the retinal ganglion cells, and various subcortical areas including the raphe nuclei, thalamic intergeniculate leaflet, and parabrachial nucleus in the brainstem (14,21,22). These subcortical nuclei initiate tau accumulation in the brain (23).

We conducted in-situ proteomics analysis on the human postmortem SCN specimen at progressive stages of AD neuropathological changes, using barcode-conjugated 57 antibodies commonly associated with dysregulation in AD. GeoMx Digital Spatial Profiling (DSP) was employed for this purpose, allowing for the quantification of spatially conserved protein expression within AVP+ and VIP+ neurons (24). As regional controls, we investigated neighboring anterior hypothalamic nuclei, such as the paraventricular nucleus (PVN) and supraoptic nucleus (SON), which also contain AVP+ neurons but demonstrate greater resilience to AD. The differentially expressed proteins (DEPs) identified through digital analysis were further validated via neuropathological and histological examination.

## Materials and methods

### Participants and neuropathological diagnosis

Participants were recruited from the Neurodegenerative Disease Brain Bank (NDBB) from Memory and Aging Center of the University of California, San Francisco (UCSF) (San Francisco, USA), until 2023, and Brazilian BioBank for Aging Studies (BBAS) from the university of São Paulo Medical School (USP) (São Paulo, Brazil), until 2023 (Fig. S1). BBAS are population-based and host a high percentage of control subjects who are not available in NDBB (25). Table 1 indicates demographic information of the participants. A total of 11 patients with amnestic AD and 7 patients with atypical AD (Braak VI), 30 cognitively normal and neuropathologically confirmed individuals (12 Braak 0, 12 Braak I, and 6 Braak II). All neurological and neuropathological assessments were performed using standardized protocols and followed the internationally accepted criteria for neurodegenerative diseases (26,27). Subjects with any other significant neurodegenerative or cerebrovascular changes were excluded from the research.

**Table 1.** Demographic characteristics of participants.

|  | Braak 0 | Braak I | Braak II | Braak VI<br>(Amnestic<br>AD) |
| --- | --- | --- | --- | --- |
| Total | 12 | 12 | 6 | 11 |
| Female, n (%) | 3 (25%) | 5 (42%) | 3 (50%) | 3 (27%) |
| Male, n (%) | 9 (75%) | 7 (58%) | 3 (50%) | 8 (73%) |
| Age of death, mean, yr (SD) | 54 (3) | 58 (6) | 71 (8) | 69 (8) |
| Disease onset, mean, yr (SD) | na | na | na | 59 (9) |

### Postmortem specimen processing

Specimen processing was followed by the UCSF NDBB and USP BBAS Neuropathological assessment protocol with NIA-AA guidelines to assess the staging of AD neuropathologic changes (REF). The Institutional Review Boards of both participating institutions approved the study. Human hypothalamus was collected from the defined area bounded by the third ventricle, optic chiasm, anterior commissure, and optic tract (Fig. 1). Specimens fixated in 10% neutral buffered formalin (how long?), and was processed and embedded in paraffin, cut into 6um thick sections, and mounted on glass slides. Region-of-interests (ROIs) were identified using Allen Brain Atlas (Allen Institute for Brain Science), human hypothalamus atlas (28), and AVP/VIP RNA probe expression using fluorescent In-situ hybridization (FISH). Fig. 1 illustrates the experimental workflow.

**Figure 1.**
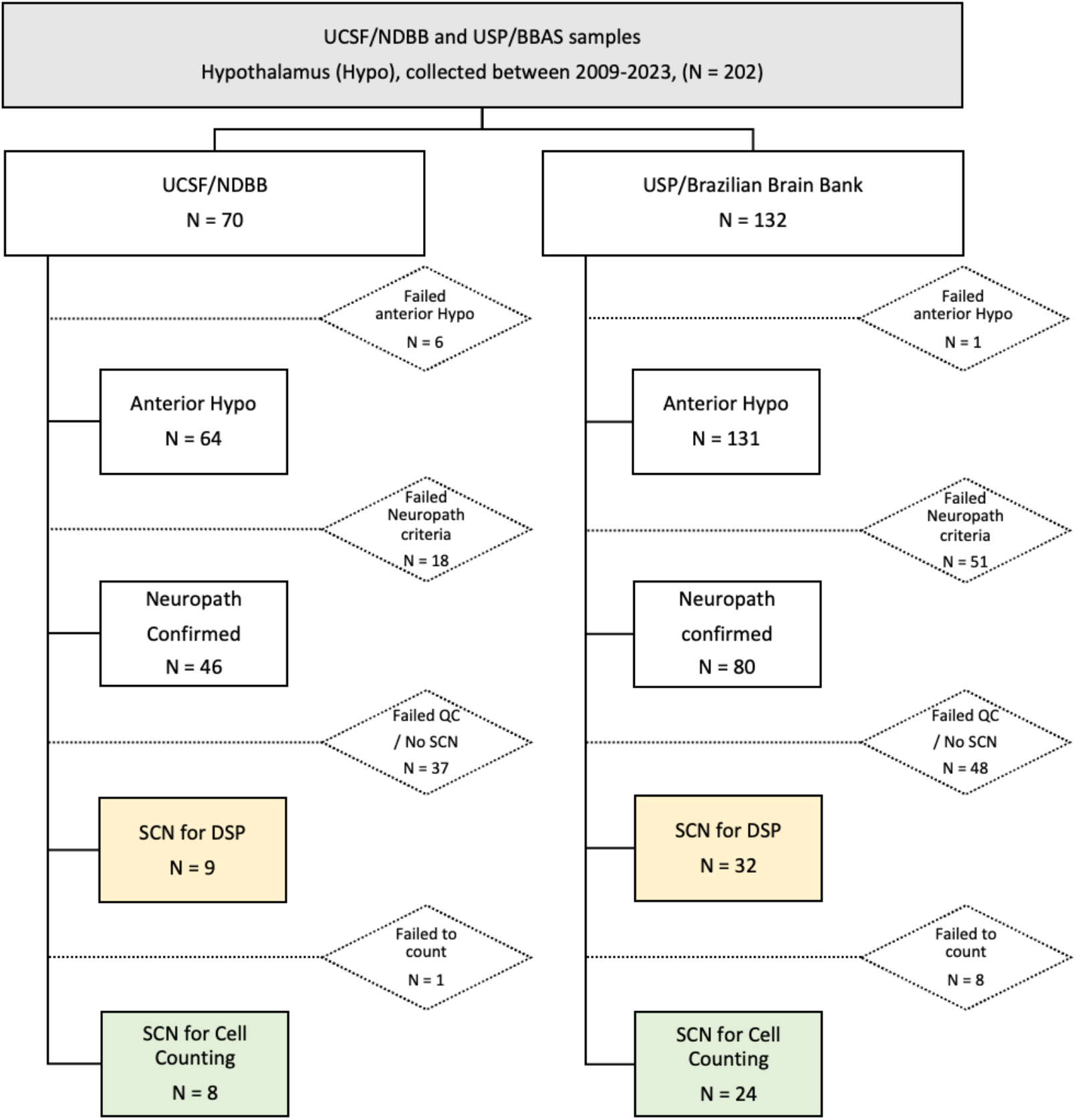
Subject ascertainment flow chart. Postmortem specimens sourced by the University California of San Francisco – Neurodegenerative Disease Brain Bank (UCSF/NDBB) and University São Paulo - Brazilian BioBank for Aging Studies (USP/BBAS) between 2009-2023. Total 32 suprachiasmatic nuclei were selected by neuropathological, anatomical, molecular criteria for the research. DSP: digital spatial profiler, Hypo: hypothalamus, N: population, Neuropath: neuropathology, QC: quality control, SCN: suprachiasmatic nucleus.

**Figure 2.**
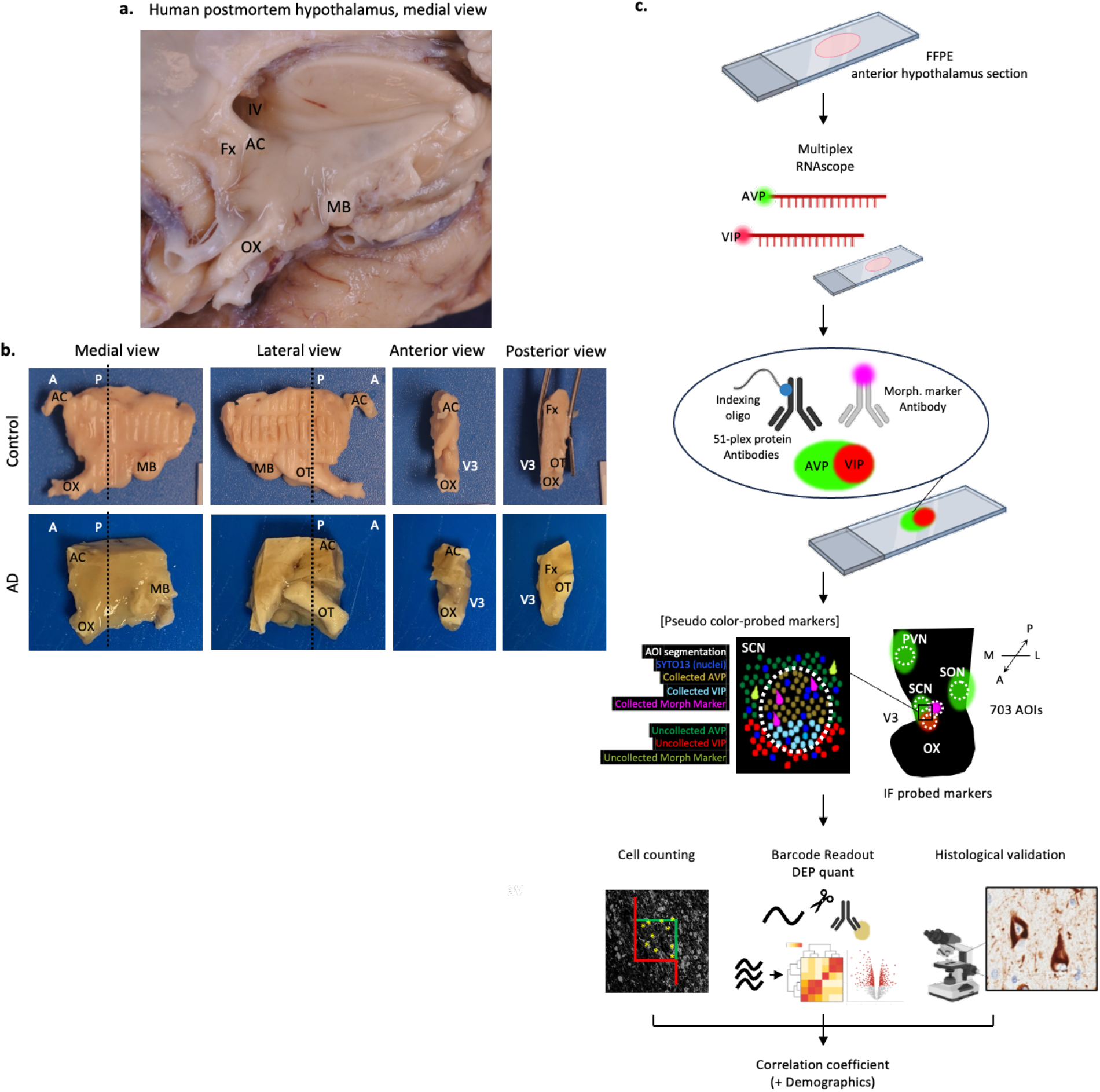
Gross anatomy of the hypothalamus and the adjoining structures, and experimental workflow. (a) Median sagittal section of the brain depicting the hypothalamus and the neighboring subcortical structures. (b) Formalin fixed hypothalamic slab in different anatomical plan. Medial, lateral, anterior, and posterior view. Posterior view is the surface of the section indicated by block dotted lines. (c) (From top to bottom) Formalin fixed paraffin embedded anterior hypothalamus sections. AVP and VIP RNA probes were hybridized by RNAscope protocol. Primary antibody mix (barcoded panel proteins and morphological markers) incubated by GeoMx DSP Protein Slide Preparation protocol. Image scanned, select region of interests (ROIs) and segment ROIs by adjusting fluorescence threshold of each channel by GeoMx DSP instrument. Photocleaved oligo barcodes were hybridized to Hyb Codes and readout by nCounter. Exported raw data was scaled by ROI size using GeoMx DSP software. The size scaled data were run background QC, normalized, batch control by bioinformatician-created R package, ‘DSPview (ver. 0.9.3)’. Fold change heatmaps were created by DSPview. The scanned images were imported to Stereo Investigator to quantify AVP and VIP population, area, and density. Protein markers were validated using immunohistochemistry in the next section of the DSP sections. A: anterior, AC: anterior commissure, AD: Alzheimer’s disease, AOI: area-of-interest, AVP: arginine vasopressin, DEP: differentially expressed protein, DSP: digital spatial profiler, Fx: fornix, MB: mamillary body, IF: immunofluorescence, OT: optic tract, OX: optic chiasm, P: posterior, PVN: paraventricular nucleus in hypothalamus, ROI: region-of-interest, SCN: suprachiasmatic nucleus, SON: supraoptic nucleus, V3: third ventricle, VIP: vasoactive intestinal peptide.

### Multiplex fluorescent In-situ hybridization (FISH) combined with immunofluorescence

Specimen sections were deparaffinized in xylene for 30min, washed in 100%, 95%, and 80% ethanol. To run the FISH assay, followed the instruction in the user manuals Formalin-Fixed Paraffin-Embedded (FFPE) Sample Preparation and Pretreatment and the RNAscope Multiplex Fluorescent Reagent Kit v2 (Cat. No. 323100, Advanced Cell Diagnostics, Newark, CA, USA). A summarized tissue pretreatment and probe binding procedure is following. Incubate the slides in hydrogen peroxide, perform target retrieval with a standard protocol, apply protease Plus, incubate probes AVP-C2 (Cat. No. 401361-C2), VIP-C3 (Cat. No. 452751-C3). RNA signals are amplified and bound fluorophores, Opal 570 for AVP-C2 and Opal 650 for VIP-C3. Slides are incubated in Sudan Black B solution to avoid autofluorescent signals.

### Digital spatial profiling with the GeoMx Human protein panels

The slides with the AVP and VIP RNA probes were processed by the GeoMx TM Digital Spatial Profiler slide preparation protocol (NanoString Technologies, Seattle, WA, USA). After an antigen retrieval with 1x citrate buffer and a blocking step with Buffer W, the slides are incubated with GeoMx Human Protein Modules for nCounter (Alzheimer’s Pathology Panel (Cat. No. 121300109), Alzheimer’s Pathology Extended Panel (Cat. No. 121300114), Neural Cell Profiling Panel (Cat. No. 121300108), Autophagy Panel (Cat. No. 121300115), NanoString Technologies, Seattle, WA, USA), CaSR (MAb 1C12D7), metabotropic gamma-aminobutyric acid (GABA) receptors GABBR1 (Abcam, ab264069, RabMAb EPR22954-47), and GABBR2 (Abcam, ab230136, RabMAb EP2411), and Alexa Fluor 594 (AF594) (Cat. No. A20182, Thermo Fisher Scientific)-conjugated AT8 (phosphor-tau, S202, and T205) (Cat. No. MN1020, Thermo Fisher Scientific) and GFAP (Cat. No.NBP1-05197AF594, Novus Biologicals) antibody. The slides were treated by 4% paraformaldehyde for a post-fixation and were stained by SYTO 13 (Cat. No. S7575, Thermo Fisher Scientific) for nuclei staining.

The slides were scanned using the GeoMx Digital Spatial Profiler (DSP) instrument at 20x magnification. The morphology of the hypothalamus nuclei (SCN, PVN, and SON) were visualized using FISH-labeled AVP and VIP probes. NFTs were labeled with AF594-conjugated AT8 antibody. Area-of-interest were selected using freeform segmentation with a size, maximum 660 µM^2^. Segmentation was performed based on human hypothalamus anatomy and physiological markers and morphological characteristics of each nucleus. Multiple sections (2∼4 sections per case) were analyzed for an unbiased DSP analysis. Total 703 of AOIs were collected.

### GeoMx DSP protein nCounter quantification

The oligonucleotide tags corresponding to bound antibodies within each cell compartment were released by UV (385 nm wavelength) photocleavage. The released oligonucleotides were recovered and dispensed into 96-well plates. The indexing oligonucleotides were dried down and resuspended in diethyl pyrocarbonate (DEPC)-treated water, hybridized to 4-color, 6-spot optical barcodes, and digitally counted using the nCounter system (NanoString Technologies, Seattle, WA, USA).

### Differentially expressed protein analysis of human GeoMx DSP data and statistics

Raw data was exported using GeoMx DSP software to normalize the digital counts using internal spike-in controls (ERCCs) and scale each AOIs to area. All batch control, post-normalization, statistical analysis, and differentially expressed protein (DEP) data visualization were performed using R software (ver. 4.3.1, R Foundation for Statistical Computing, Vienna, Austria, https://www.R-project.org/).

### Histological examination and statistics

The number of cells and the size of ROIs were estimated at a series of sections spanning the whole SCN using Stereo Investigator probe (ver. 2023, MBF Bioscience LLC, Williston, VT, USA). The scanned images of histological slides that underwent GeoMX DSP analysis were loaded into the Stereo Investigator software and were used to quantify the number of AVP+ and VIP+ neurons. Two Braak 0 and two Braak I cases used for DSP analysis were excluded from the neuronal quantification because the tissue physically damaged or the ROI was not available entirely, a prerequisite for quantification. Image magnification was corrected by the embedded scale bar if the exported DSP images have different scale. Insoluble tau population was quantified using semi-quantification methods. The same slides performed DSP analysis were re-scanned using Axioscan 7 (Carl Zeiss AG, Oberkochen, Germany) to quantify the insoluble tau. Data consistency in counts were confirmed between two trained, blinded investigators.

## Results

### Neuromodulatory neurons in the suprachiasmatic nucleus progressively decrease from Braak stage II onwards

The alterations of SCN neurons are illustrated in Figure 3. SCN is in the superior side of the optic chiasm and medial anterior hypothalamus next to the third ventricle probed with AVP (green) and VIP (red) (Fig. 3a). AVP+ neurons are present in all three nuclei, i.e., SCN, PVN and SON (Fig. 3a). Number of AVP+ and VIP+ neurons decreased in AD progressive stages, classified as Braak stages (Fig 3b and c; estimated loss of 0.1 per Braak stage, P < 0.005). The neuronal density remained the same across progressive Braak stages due to decreased area expressing AVP+ and VIP+ neurons (Fig, 2d-g, P = 0.671). Braak stage VI showed a 50% reduction in the count of SCN neurons, with 50% (AVP) and 30% (VIP) reduction in SCN area compared to Braak stage 0 cases. Interestingly, the decrease found at Braak stage II in both neurons they were already ∼40% smaller than in Braak stage 0 cases, showing the AD-related neuromodulatory changes in SCN precede the severe cortical changes in AD (see a loess regression, black dotted line, Fig. 3b-e). This finding was evenly found regardless of sex and age of death (Data were corrected by sex and age of death). Further, AVP+ neurons and their expressing area in the PVN and SON remained intact compared to the neurons in the SCN (Fig. S2).

**Figure 3.**
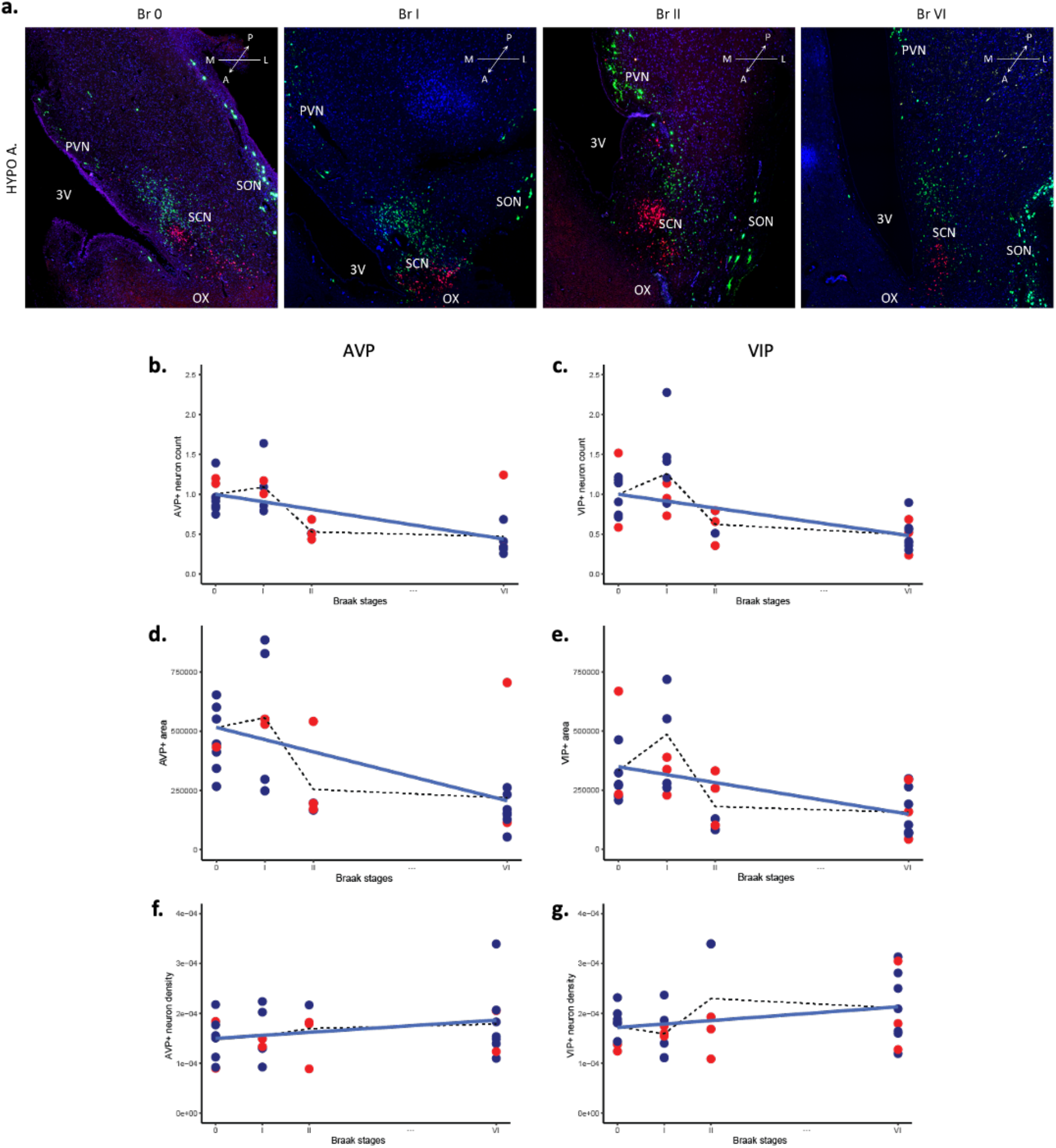
Association between histological quantification and progressive Braak stages in the suprachiasmatic nucleus. (a) Three anterior hypothalamic nuclei were probed for each Braak stage with AVP and VIP RNA. Scale bar: 1mm. (b-g) Linear regression models indicate a negative correlation between the Braak stage and neuron population, as estimated by the Stereo Investigator, contoured nucleus size, and density (population/size). (b) AVP+ and (c) VIP+ neuron populations estimated by Stereo Investigator in SCN. Data was normalized by ratio to Braak stage 0 (β = X with P < 0.005 and 95% CI: X). (d) AVP+ and (e) VIP+ region size (β = X with P < X and 95% CI: X). (f) AVP+ and (g) VIP+ neuron density (β = X with P = 0.671 and 95% CI: X). Individuals were color-coded as male (blue) and female (red). Sex and age of death were corrected. The block dotted lines connect the mean value of each Braak stage. A: anterior, AVP: arginine vasopressin, DEP: differentially expressed protein, DSP: digital spatial profiler, Fx: fornix, IF: immunofluorescence, L: lateral, M: medial, OX: optic chiasm, P: posterior, PVN: paraventricular nucleus in hypothalamus, SCN: suprachiasmatic nucleus, SON: supraoptic nucleus, V3: third ventricle, VIP: vasoactive intestinal peptide.

### Suprachiasmatic nucleus is more susceptible to accumulate phospho-tau than amyloid-ß

Elevated pathological tau accumulation in the SCN was confirmed by DSP and histological examination (Fig. 4). Tau fibril aggregation and neurofibrillary tangle detected by p-tau immunoreactivity were found at Braak stage II (Fig. 4a). The pathological tau forms were significantly increased at Braak stage II. They became exponentially abundant at Braak stage VI compared to Braak stage 0 (Fig. 4b). DSP result confirmed that SCN exhibited significant p-tau up-regulation (S404) at Braak stage II and continued to escalate at Braak stage VI (Fig. 4c). Three representative p-tau species, i.e., p-S404, p-S396 and p-S214, were identified in the SCN in both AVP+ and VIP+ neurons and significantly upregulated in Braak stage VI compared to other earlier Braak stages (Fig. 4g, j, m). Contrary to SCN, the PVN and SON did not show any substantial increase in the level of p-tau in AVP+ neurons in the neighboring nuclei, PVN and SON compared to Braak stage 0 (Fig. 4h, k, n for PVN, Fig. 4f, i, l, o for SON, and Fig. S4). The up-regulated p-tau proteins were validated by p-tau immunoreactivity in the adjacent DSP sections. Tau fibrils and neurofibrillary tangles were predominantly examined in the SCN and surrounding the SCN but rarely in the neighboring nuclei, such as the PVN and SON (Fig. 4q).

**Figure 4.**
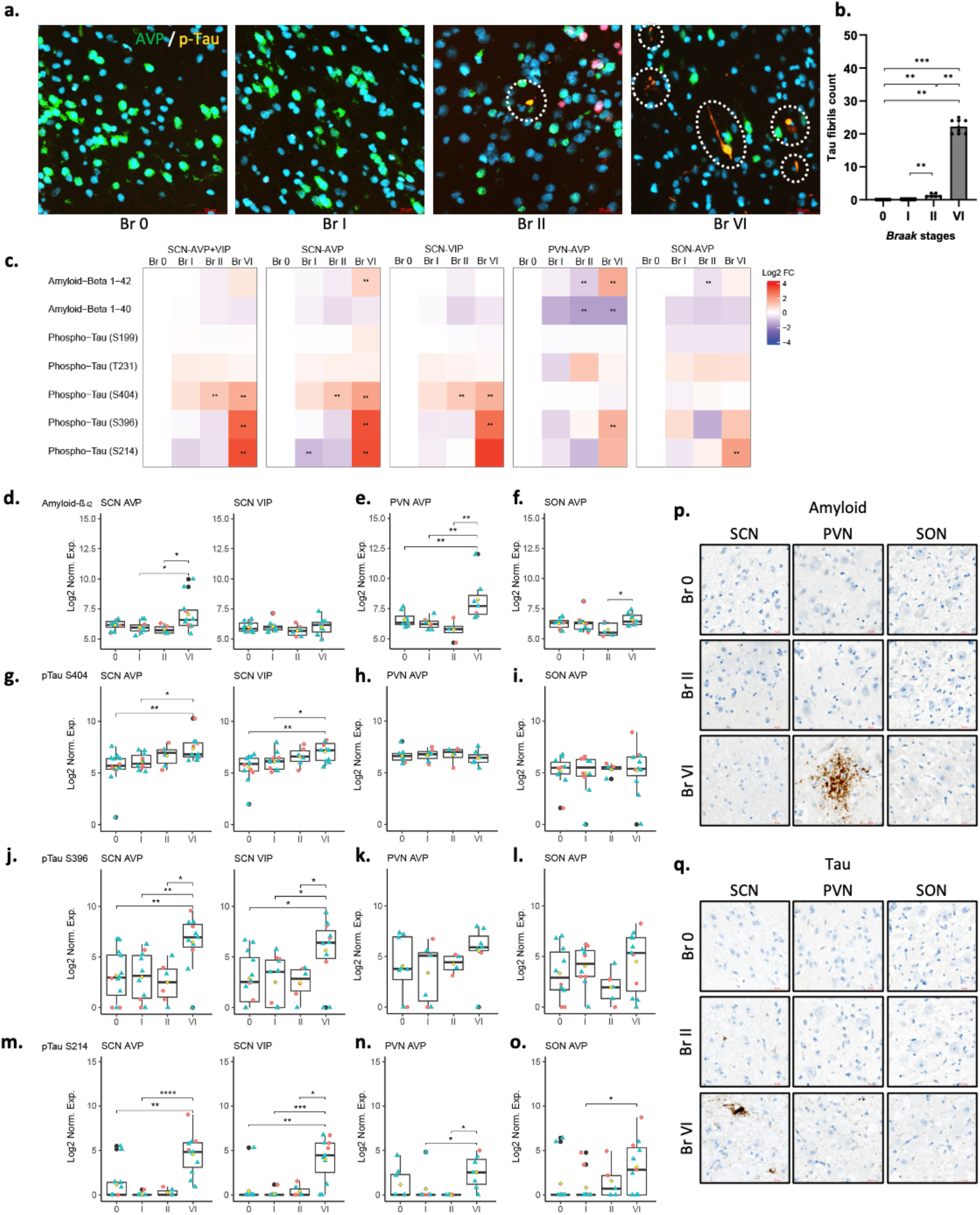
Amyloid and tau pathology in suprachiasmatic nucleus in progressive Braak stages. (a) Tau fibril immunoreactivity (AT8) in the SCN in each Braak stage. Scale bar: 50um. (b) Comparison of tau fibril count in the SCN in each Braak stage. Individuals were coded block dots. P-values were analyzed by *Wilcoxon* test (**; P < 0.01, ***; P < 0.001, Holm-adjusted). (c) Comparison between differentially expressed proteins (DEP) using fold change heatmap. Each column displays DEP in each Braak stage compared to Braak stage 0. Log2 Fold-change values are color-annotated, with red (up-regulated proteins) and blue (down-regulated proteins). Each row shows pathoproteins quantified by the barcode. P-values were calculated using pseudobulk analysis including permutation test (**; P < 0.01). (d-o) Multiple comparison of pathoprotein expression in each Braak stage by *Wilcoxon* test (*; P < 0.05, **; P < 0.01, ***; P < 0.001, Holm-adjusted by Braak stages). Individuals were color/shape coded as Male (blue/triangle) and Female (red/circle). Sex and age of death were corrected. Log2 Norm. Exp.: log2 of normalized expression. (d, g, j, m) SCN AVP+ (left) VIP+ (right) neuron. (e, h, k, n) PVN neuron. (f, i, l, o) SON neuron. (d-f) amyloid-β 1-42. (g-i) amyloid-β 1-40. (j-l) p-tau S396. (m-o) p-tau S214. (p) amyloid-β 1-42 immunoreactivity in each nucleus and Braak stages. Scale bar: 50um. (q) PHF1 immunoreactivity in each nucleus and Braak stages. Scale bar: 50um.

In contrast to the notable tau pathology, amyloid-ß 1-42 levels showed a substantial increase in the SCN at only Braak VI. This elevation in amyloid-ß 1-42 level was restricted to AVP+ neurons in the SCN (Fig. 4c, d). Furthermore, the amyloid-ß 1-40 level did not show any change in the SCN (Fig. 4c and Fig. S4a). In addition to the SCN, SON AVP+ neurons did not show amyloid pathology (Fig. 4f and Fig. S4c). Interestingly, PVN AVP+ neurons increased amyloid-ß 1-42 and decreased 1-40 expression in severe AD (Fig. 3e and Fig. S4b), suggesting a significantly higher amyloid-ß ratio 42 to 40 in Braak stage VI. Further, amyloid-ß 1-42 immunoreactivity confirmed no plaque in the SCN (Fig. 4p). However, PVN showed neuritic plaques (Fig. 4p). In contrast, SON showed tiny dotty aggregations within a cell (Fig. 4p). Given to the AD pathology profiling, SCN exhibited primarily tau selective vulnerability compared to other neighboring nuclei. PVN showed mild tau and amyloid pathology, and SON bears very mild tau pathology but rarely amyloid.

### Glial proteins are dysregulated in the SCN neurons before the advent of p-tau pathology

Astrocytes are abundant in the SCN, with high endogenous expression level of GFAP and S100B in both AVP+ and VIP+ regions of the SCN in Braak stage 0 (Fig. S5a). We observed higher expression of astroglia markers than other types of glial markers such as microglia and oligodendrocytes (Fig. S5a). Immunoreactivity for GFAP confirmed that SCN contain dense astrocyte population (data not shown). We found that the astrocytic markers were up-regulated even in Braak stage I, even before p-tau accumulated (Fig. 5b, c). Multiple comparison analysis revealed a selective vulnerability, showing the more significance in SCN-VIP+ than SCN-AVP+ neurons (Fig. 5e,h,k). The astrocyte markers showed endogenous high expression PVN and SON at Braak stage 0 (Fig. S5b) and later stages (Fig. 5d, e), excluding Braak stage I in PVN (Fig. 5d). Intriguingly, GFAP levels in the PVN at Braak stage I were significantly low in the DSP analysis, but multiple comparison analysis showed sustained level (Fig. 5i). The upregulated astroglia proteins in the early stages had a similar expression level at Braak stage VI to the level at Braak stage 0. Contrary to astrocytic markers, oligodendrocyte did not show any change in the SCN.

**Figure 5.**
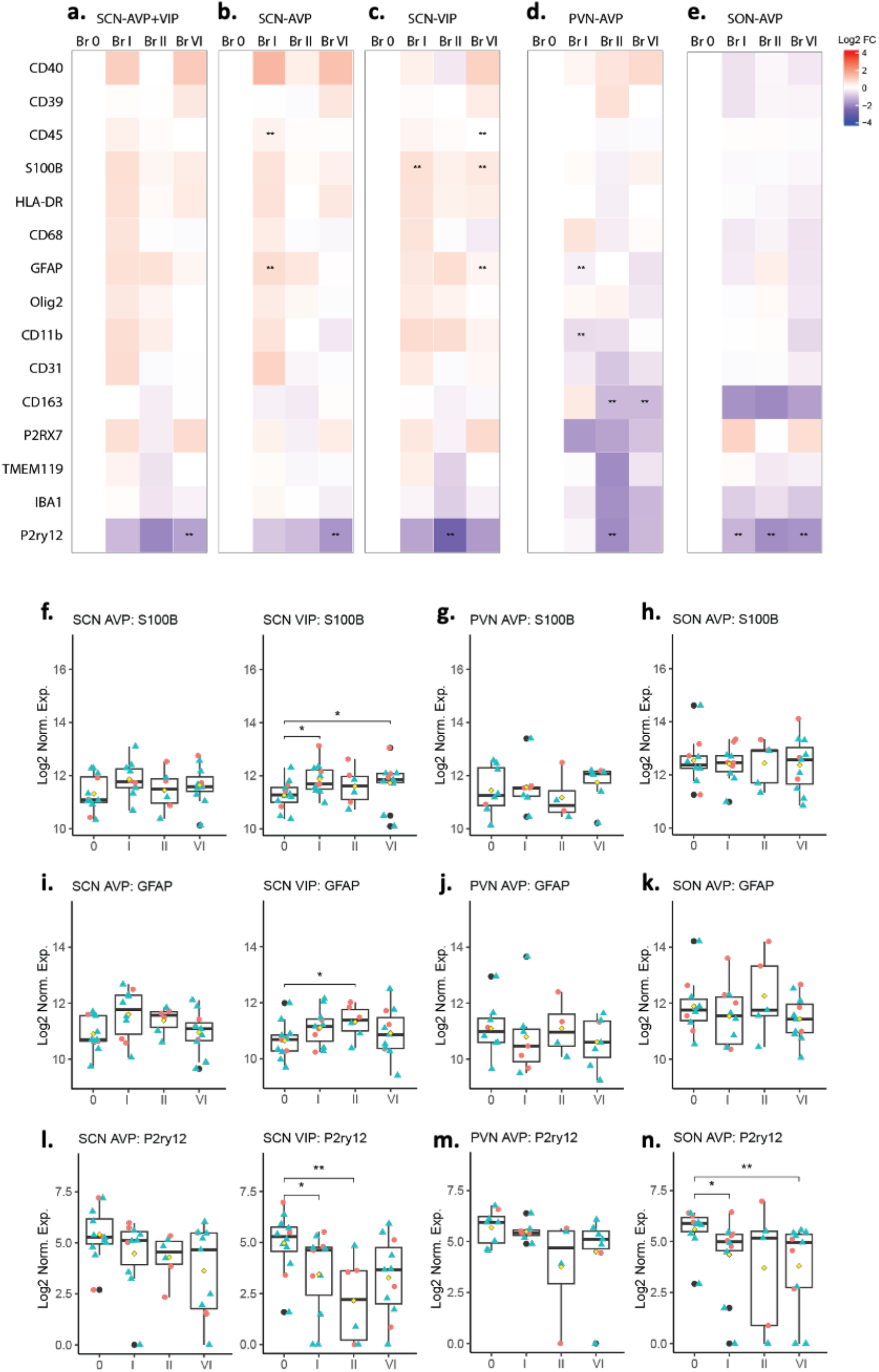
Glial markers in the suprachiasmatic nucleus in progressive Braak stages. (a-e) Comparison between differentially expressed proteins (DEP) using fold change heatmap. Each column displays DEP in each Braak stage compared to Braak stage 0. Log2 Fold-change values are color-annotated, with red (up-regulated proteins) and blue (down-regulated proteins). Each row shows pathoproteins quantified by the barcode. P-values were calculated using pseudobulk analysis including permutation test (**; P < 0.01). (a) Combined SCN AVP+VIP+ neuron. (b) SCN AVP+ neuron. (c) SCN VIP+ neuron. (d) PVN neuron. (e) SON neuron. (f-n) Multiple comparison of pathoprotein expression in each Braak stage by *Wilcoxon* test (*; P < 0.05, **; P < 0.01, Holm-adjusted by Braak stages). Individuals were color/shape coded as Male (blue/triangle) and Female (red/circle). Sex and age of death were corrected. Log2 Norm. Exp.: log2 of normalized expression. (f, i, l) SCN AVP+ (left) VIP+ (right) neuron. (g, j, m) PVN neuron. (h, k, n) SON neuron. (f-h) S100B. (i-k) GFAP. (l-n) P2ry12.

Microglial related proteins showed dysregulation throughout the progression of AD, especially phagocytic microglia. The P2RY12, a homeostatic microglial marker involved in misfolded protein phagocytosis, decreased over Braak stages, and demonstrated differential pattern in areas adjacent to SCN-AVP and SCN-VIP neurons (Fig. 5b, c). Multiple comparison valued the decrease was significant at Braak stage I (Fig. 5k). In addition to the SCN, altered the P2RY12+ protein co-localized to the SON neuron and demonstrated significant low levels even at Braak stage I compared to the Braak 0 (Fig. 5m).

### SCN neuropathology demonstrates selective vulnerability SCN neuropathology demonstrates selective vulnerability in AD variants

SCN analyses revealed significant pathological change in AD subjects. We investigated whether different AD variants exhibit the same propensity of molecular changes in the SCN (Fig. 6). Total seven SCN investigated from atypical AD (Atyp.AD) subjects compared to twelve Braak 0 and eleven amnestic AD (Am.AD) (Fig. 6a). Interestingly, the SCN in Atyp.AD showed more resilience to tau fibril accumulation (Fig. 6b, c). Same as Am.AD, amyloid-ß immunoreactivity confirmed that amyloid-derived plaque was not found in the SCN (data not shown). DSP analyses demonstrated that the levels of other proteins show a different expression pattern in Am.AD vs. Atyp.AD (Fig. 6d). In figure 6d, Am.AD exhibited a more dynamic pathoprotein expression in the SCN than Atyp.AD. In addition to p-tau proteins, neprilysin, a metalloprotease that degrades Aß plaques, indicating compensatory mechanism in the SCN. Intriguingly, higher expression of HSC70, GBA, and TFEB is associated with chaperone-mediated autophagy due to oxidative stress or pathological conditions with misfolding of cytosolic proteins, in response to increased p-tau. Increased PSEN1 in Am.AD can explain compensation for malfunctioning lysosomes rather than an enzymatic activity for Aß cleavage. None of these proteins are highly expressed in Atyp.AD. Moreover, the profound change in MAP2 levels suggests activation of protective pathways in SCN in Atyp.AD.

**Figure 6.**
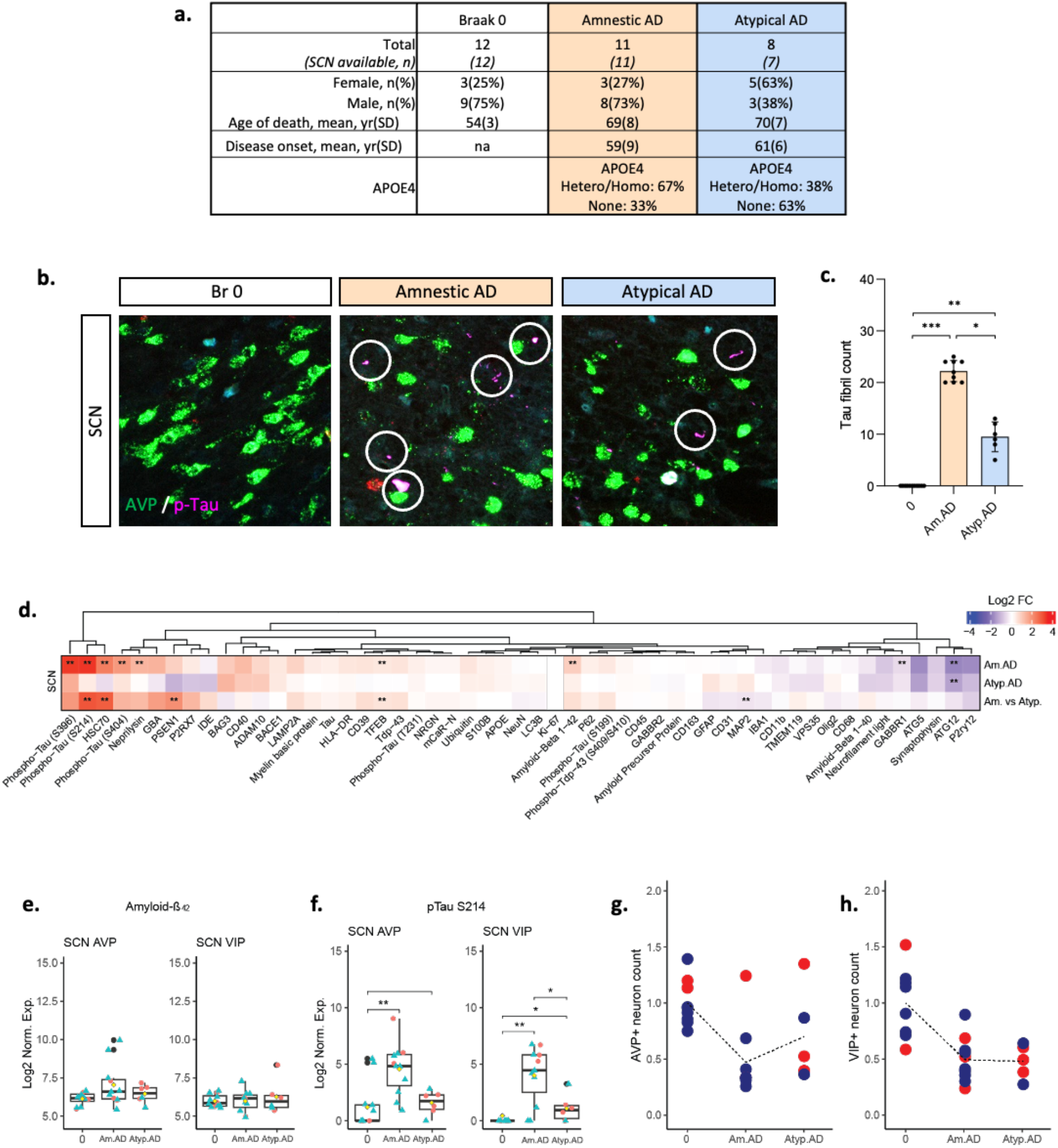
Comparison of pathoprotein expression and histological change between Alzheimer’s disease variants. (a) Summary of subject information for AD variants investigation. (b) Tau fibril immunoreactivity (AT8) in the SCN in each Braak stage. Scale bar: 50um. (c) Comparison of tau fibril count in the SCN in Braak stage 0, amnestic AD (Am.AD) and atypical AD (Atyp.AD). Individuals were coded block dots. P-values were analyzed by *Wilcoxon* test (*; P < 0.05, **; P < 0.01, ***; P < 0.001, Holm-adjusted). (d) Comparison between differentially expressed proteins (DEP) using fold change heatmap. First and second row displays DEP in each Am.AD and Atyp.AD compared to Braak stage 0. Last row displays DEP in Am.AD compared to Atyp.AD. Log2 Fold-change values are color-annotated, with red (up-regulated proteins) and blue (down-regulated proteins). Each column shows pathoproteins quantified by the barcode. P-values were calculated using pseudobulk analysis including permutation test (**; P < 0.01). (e, f) Multiple comparison of pathoprotein expression in Braak stage 0, Am.AD and Atyp.AD by *Wilcoxon* test (*; P < 0.05, **; P < 0.01, Holm-adjusted by Braak stages). Individuals were color/shape coded as Male (blue/triangle) and Female (red/circle). Sex and age of death were corrected. Log2 Norm. Exp.: log2 of normalized expression. (e) amyloid-β 1-42 in SCN AVP+ (left) VIP+ (right) neuron. (f) p-tau S214 in SCN AVP+ (left) VIP+ (right) neuron. (g, h) Comparison of neuron population estimated by Stereo Investigator. (g) AVP+ and (h) VIP+ neuron. Data was normalized by ratio to Braak stage 0. Individuals were color coded as Male (blue) and Female (red). Sex and age of death were corrected. The block dotted lines connect the mean value of each condition.

Neuronal population measured in each AVP+ and VIP+ SCN neurons. The number of both neurons were reduced in AD subjects compared to Braak stage 0, however, the AVP+ neurons in Atyp.AD sustained less damage than in Am.AD (Fig. 6g). However, this differential effect on the different neuromodulatory cells warrants further examination with more subjects.

## Discussion

The results presented in this paper delineate the molecular and pathological alterations identified within the human SCN throughout the progressive Braak stages. Leveraging the morphological and physiological features of the SCN, GeoMx digital spatial protein analysis was utilized to investigate SCN neuropathology. We found neuronal loss and a selectively vulnerability to tau accumulation in the SCN at Braak stage II, occurring before the severe neuropathological changes in cortical areas observed in AD. Conversely, the SCN exhibited resilience to amyloid pathology. Subsequently, we identified dysregulated astrocytes and microglia involvement preceding tau accumulation in the SCN. These findings were predominantly found in Am.AD SCN, whereas the AD variant exhibited greater resilience to AD pathology in the SCN. The neighboring nuclei, containing the same type of neuromodulatory cells as the SCN, exhibited varying degrees of selective vulnerability. The PVN was found to be less vulnerable to tau pathology but more prone to amyloid accumulation, whereas the SON exhibited lower susceptibility to AD pathology. This study emphasizes the importance of comprehending the molecular vulnerability of neurons and glial cells in the SCN, providing evidence for the circadian disorders observed in AD patients.

The etiology of AD is heterogeneous and involves multipathogenic and environmental changes (29). For this reason, the driving force behind the early susceptibility of the SCN pathology is controversial. However, there is evidence that hypothalamic nuclei and brainstem nuclei, which connect directly to the SCN and contribute to circadian-sleep-wake regulation, are vulnerable to tau-associated neurodegeneration in patients with Alzheimer’s disease (30), leading to the neuronal death was associated with clinical sleep disturbances (7). Although it is a challenging to distinguish the neuronal basis matrix of abnormal circadian behavior and physiology from the multiple pathways of the metabolism, finding a direct association between the pathological changes in the SCN and circadian physiological abnormalities remains an exploration, suggesting the development of objective examination criteria for evaluating the circadian function.

Investigating SCN neuronal-specific susceptibility is also a crucial point. Preliminary research has revealed the SCN neuronal specific susceptibility is also crucial point to investigate. Preliminary research revealed that the loss of AVP+ neurons in the SCN identified in AD patients (16,31,32). AVP+ neurons through the PVN, which received the efferent signals from the SCN promote wake-promoting nuclei (33), and the neuronal activity in the SCN regulates clock gene expression and circadian behavior in animal model studies (20,34,35). Likewise, VIP+ neurons are essential to process light-mediated circadian cue (36,37), and degenerated in AD patients (15). Therefore, it supports our ongoing study, which involves a detailed anatomical analysis of the subregion of the SCN, focusing on the of AVP+ and VIP+ neurons.

The role of astrocyte in the SCN regulating circadian genes supports the increased astrocyte marker on the SCN neurons in the progression of ADNC (38–41). Further studies are required to validate the morphology and distribution of glial cells in the SCN. Selective vulnerability exhibited differential neuropathologic alterations between the SCN, PVN, and SON, prompting questions about the potential mechanisms underlying this alteration. Additionally, it raises a significance about the extent of association between each nucleus and other circadian/sleep/wake-promoting nuclei in the SCN circuitry (14,42).

## Supplementary information

**Supplementary figure 1.**
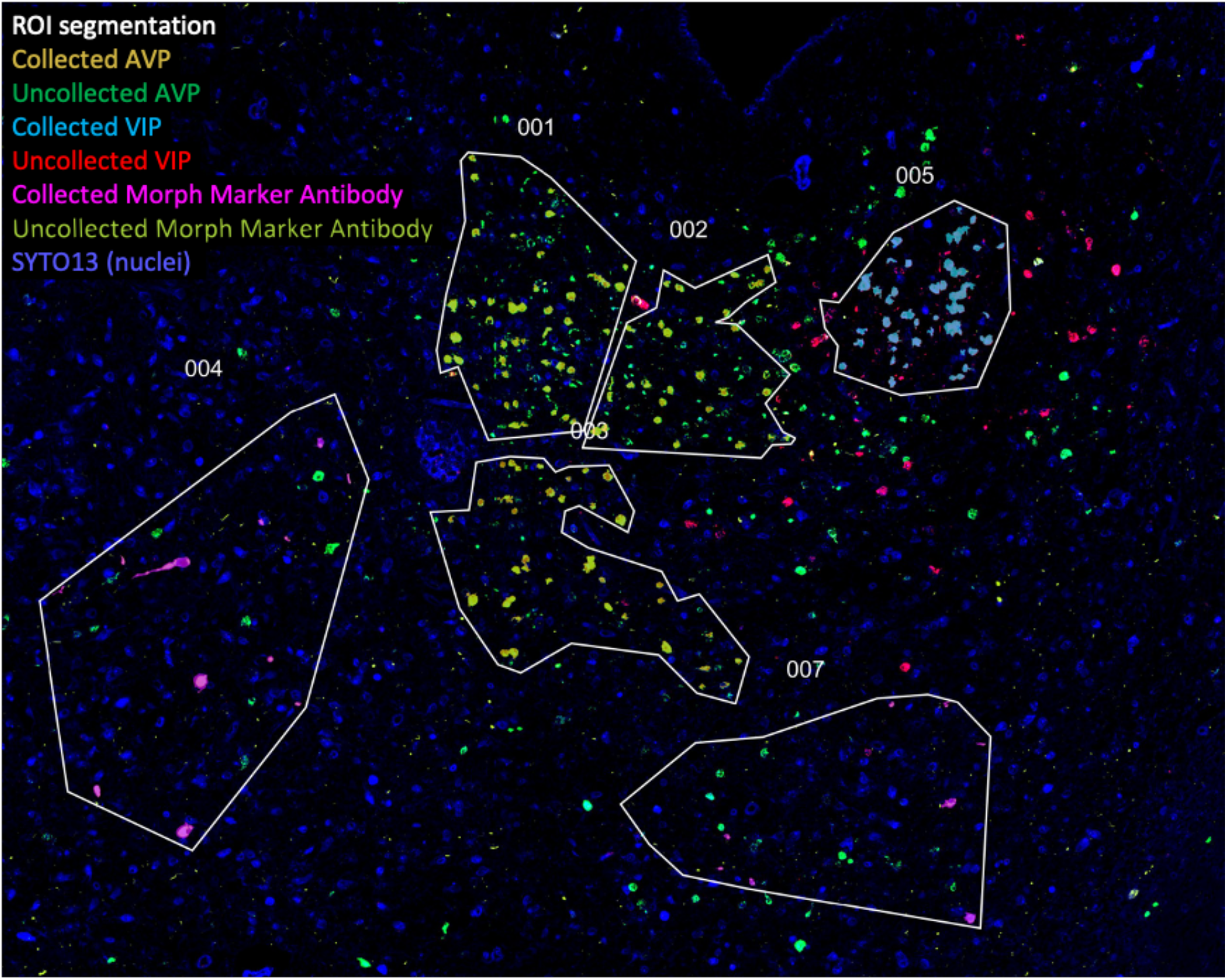
Area of interest (AOI) segmentation within region of interest (ROI) using the digital spatial profiler instrument.

**Supplementary figure 2.**
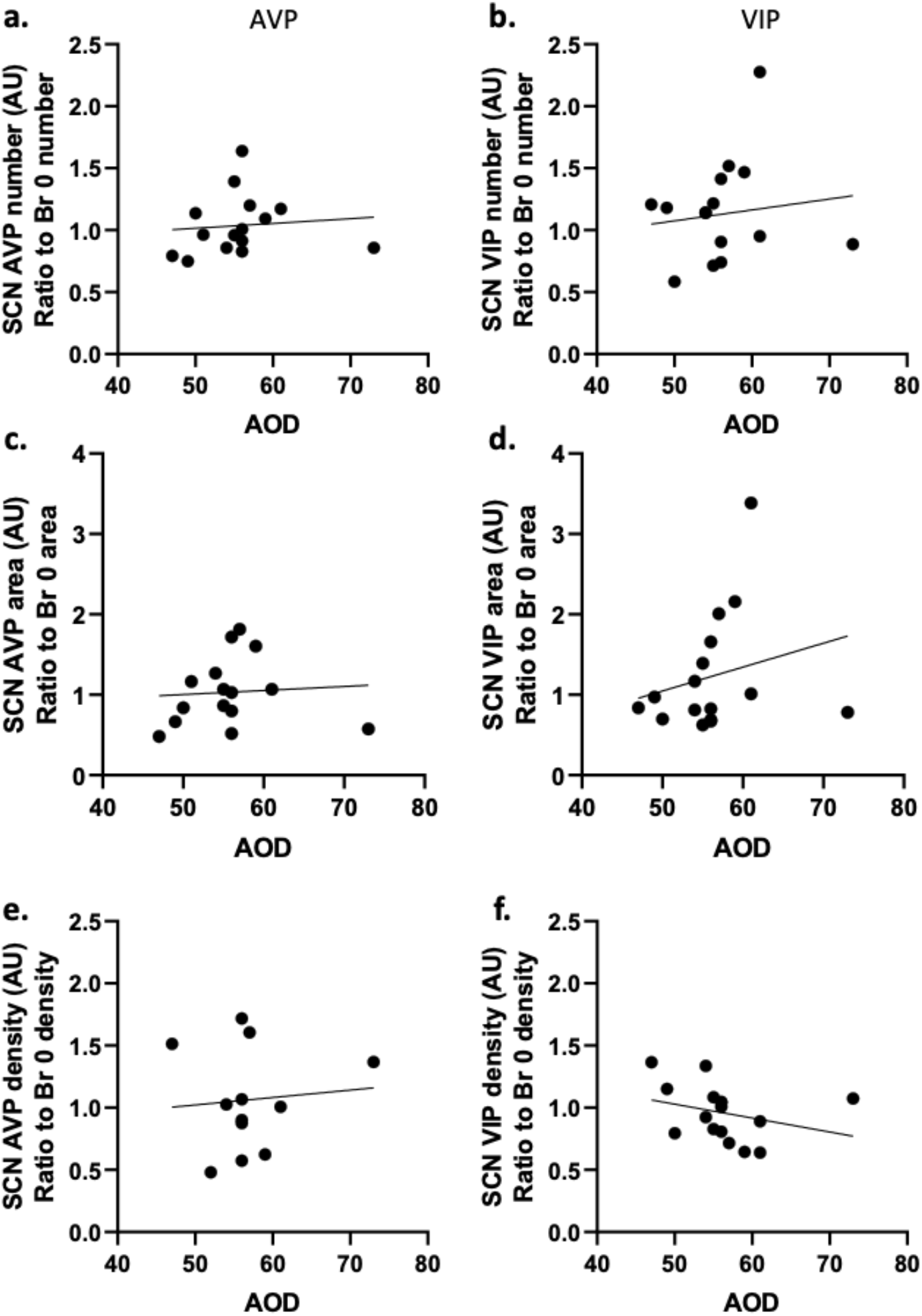
Association histological examination in the suprachiasmatic nucleus with normal aging of the subjects (Braak stage 0 and I)

**Supplementary figure 3.**
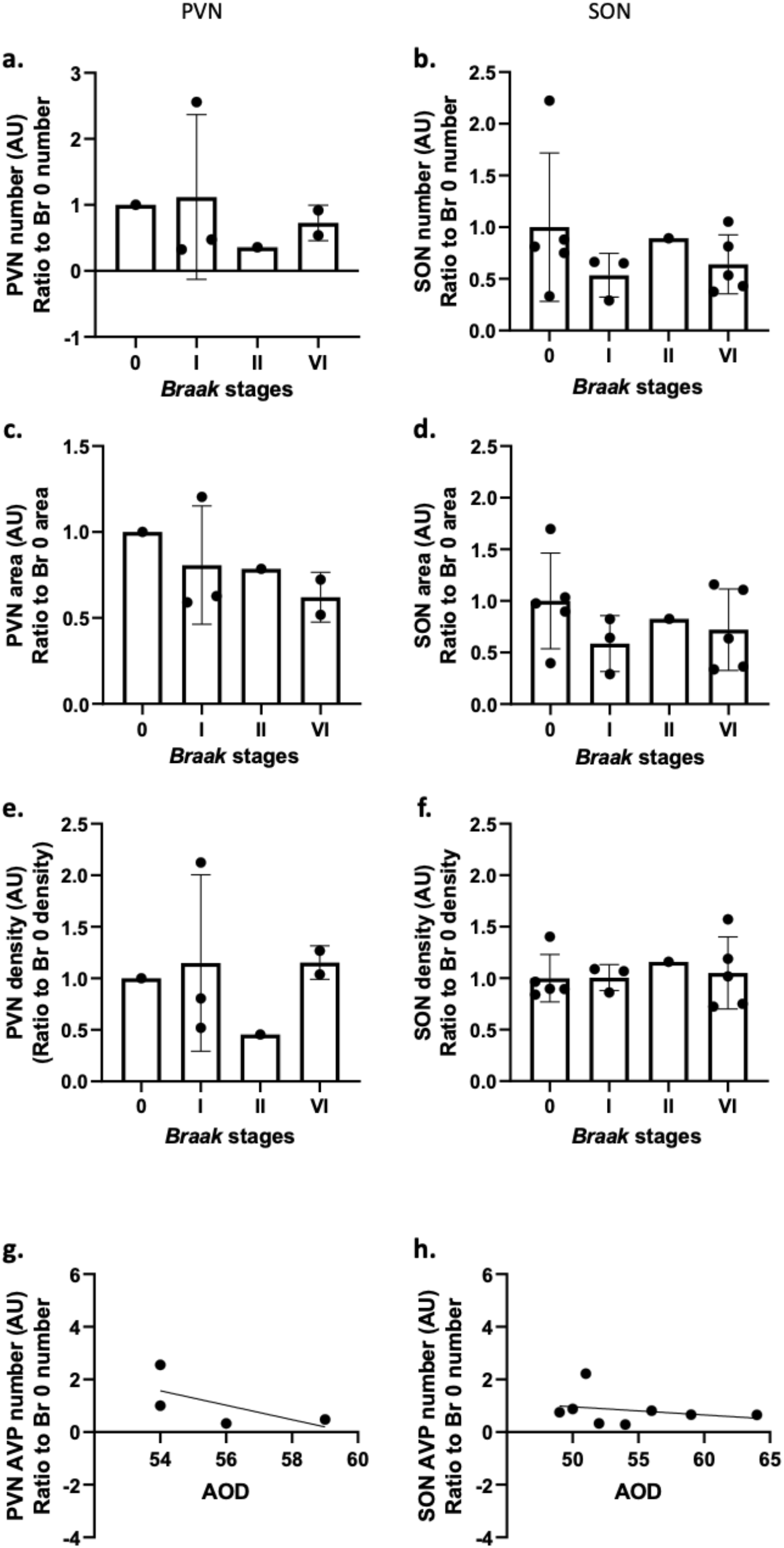
Histological examination in the paraventricular nucleus and supraoptic nucleus.

**Supplementary figure 4.**
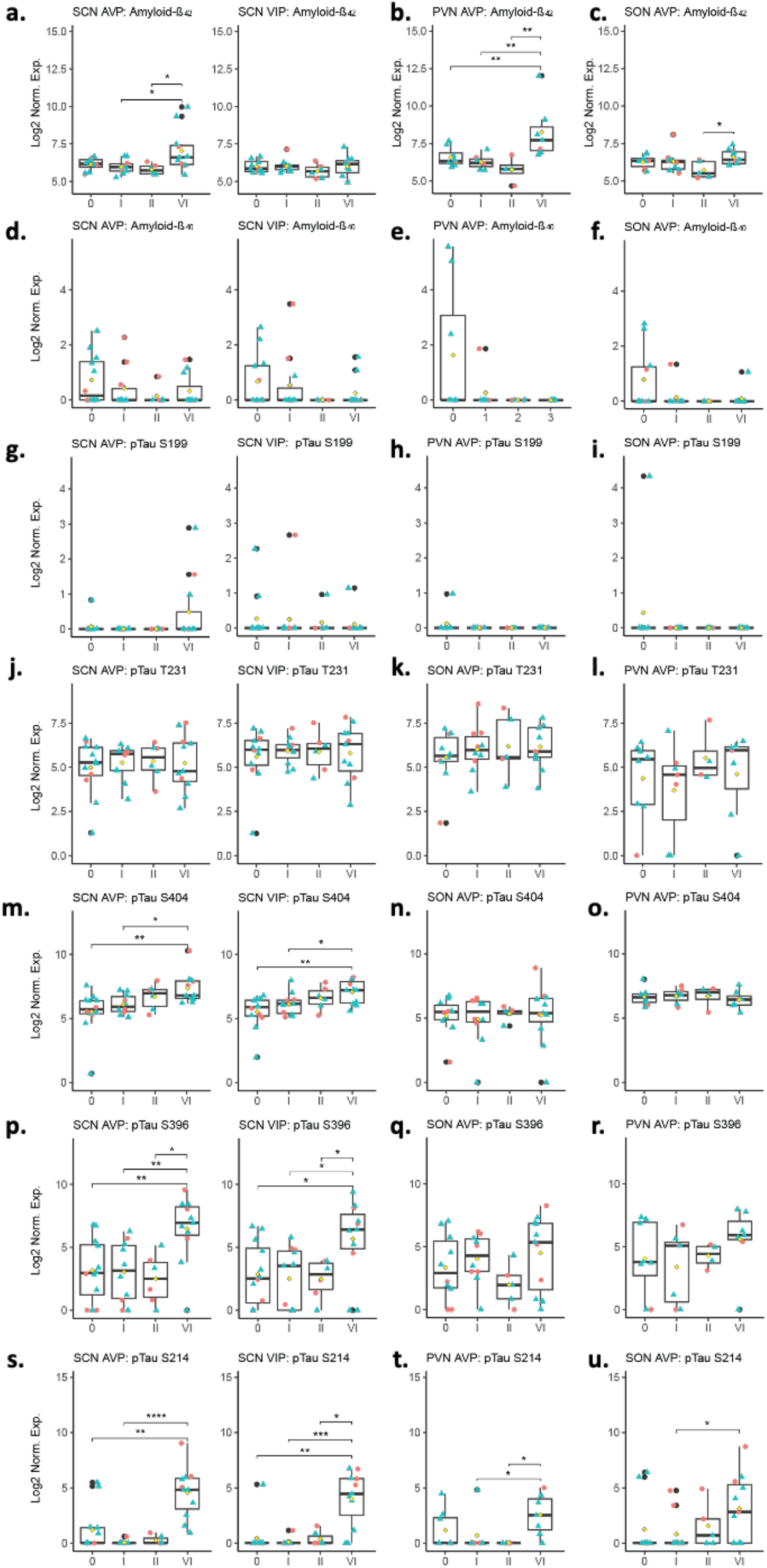
Multiple comparison analysis of differentially expressed amyloid and p-tau protein in anterior hypothalamic nuclei using digital spatial profiler (DSP)

**Supplementary figure 5.**
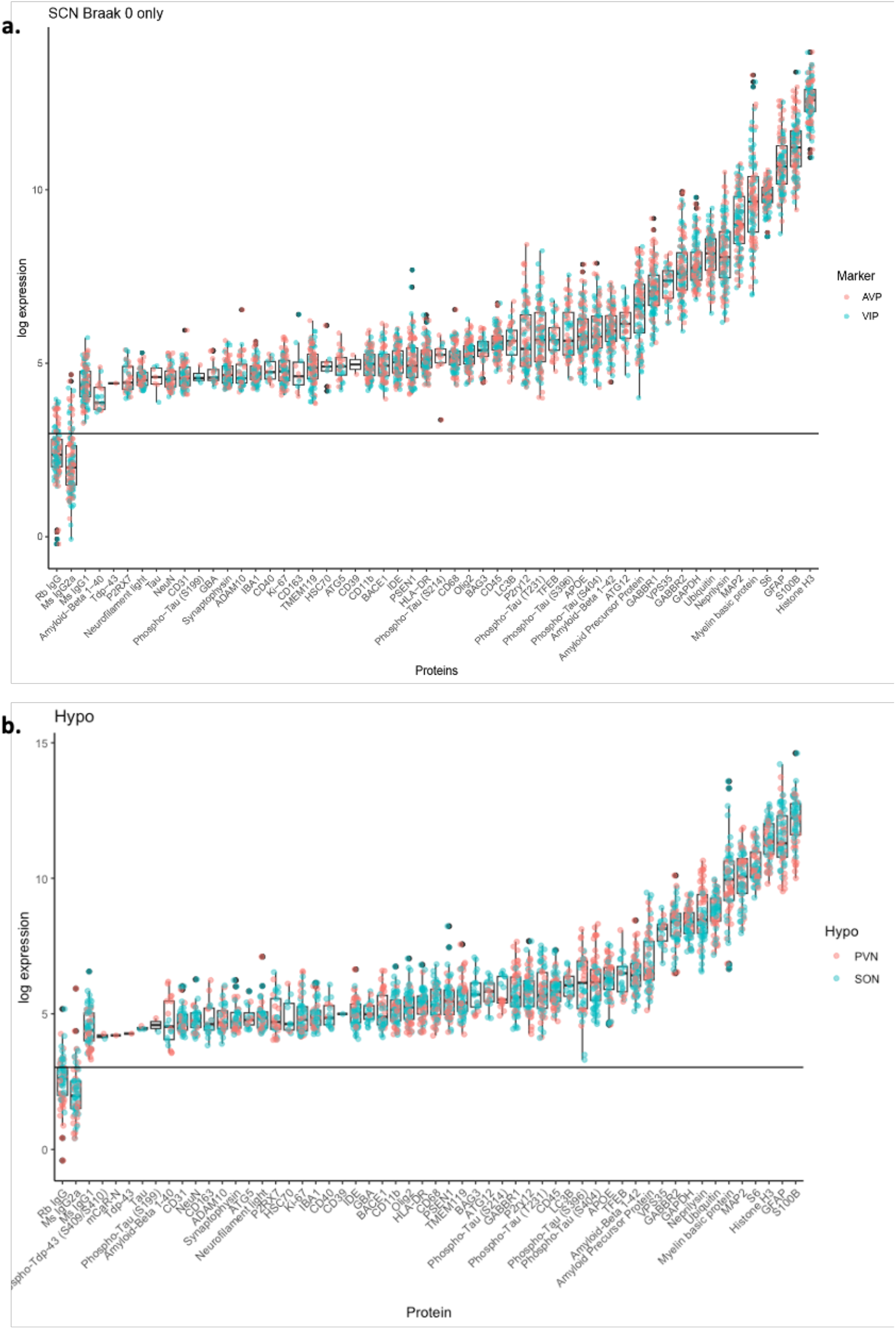
Endogenous protein expression level of digital spatial profiler (DSP) modules in the human suprachiasmatic nucleus at Braak stage 0.

## Notes

### Competing Interest Statement

The authors have declared no competing interest.

### Summary of Updates

The manuscripts were imporved with a concise introduction, clearer results accompanied by updated figures, and an enhanced causality association between histological examination, pathoprotein accumulation, and glial cell involvement. Additionally, a comparison between AD variant, amnestic AD, and control groups was included in the study.

